# Sex-dependent rewarding effects of nicotine in mice are prevented by the atypical antipsychotic aripiprazole

**DOI:** 10.64898/2026.11.18.767606

**Authors:** Rayssa C. Briânis, Lucas G. Bedeschi, Lia P. Iglesias, Samia R. L. Joca, Daniele C. Aguiar, Fabrício A. Moreira

## Abstract

Nicotine stands amongst the most commonly used drugs in contemporary societies. Preparations for nicotine consumption range from traditional tobacco products (smoking) to modern electronic cigarettes (vaping). However, its rewarding and reinforcing effects vary as a function of dose and biological sex. In addition, few treatments exists for nicotine use disorders Here we sought to characterize the effects of nicotine on anxiety in the elevated plus maze (EPM) and reward in the conditioned place preference (CPP) paradigm in male and female mice and test the hypothesis that aripiprazole, a partial dopamine agonist, prevents nicotine-induced CPP. Nicotine effects were tested at 0.25, 0.5 and 1 mg/kg. The highest dose reduced basal locomotion in both males and females exposed to the open field and reduced open arm exploration in males in the EPM. Thus, the apparent anxiogenic-like effect is confounded by a reduction in basal exploratory activity. In the CPP paradigm, nicotine 0.5 mg/kg served as a reward conditioned stimulus in males, but not females. Finally, aripiprazole (0.1, 1 and 10 mg/kg) consistently prevented the acquisition of nicotine (0.5 mg/kg) CPP in males. These findings show a sexually dimorphic effect of nicotine in mice and suggest the antipsychotic aripiprazole as a potential pharmacological approach for reducing nicotine reward.

## Introduction

Tobacco smoking remains one of the major causes of morbidity and mortality globally, contributing to cardiovascular and respiratory diseases in addition to the development of several types of cancer (1). Despite reductions in conventional cigarette consumption, electronic nicotine delivery systems (“vapes”) have renewed concern regarding nicotine addiction, particularly among adolescents and young adults (2). Nicotine is the main psychoactive compound responsible for tobacco addictive effects and is well known for its strong association with environmental cues that sustain drug-seeking behavior and relapse (3). Indeed, cue-induced craving represents a major challenge during abstinence, as exposure to contexts previously associated with nicotine use can trigger compulsive seeking even after prolonged cessation increasing vulnerability to relapse (3). From a neurobiological perspective, nicotine exerts its effects primarily through nicotinic acetylcholine receptors (nAChRs) located on dopaminergic neurons in the ventral tegmental area (VTA), increasing dopamine release in the nucleus accumbens (NAc), a key component of the brain reward circuit (3,4). Nicotine also modulates GABAergic neurons projecting from the VTA to the amygdala (Amg) mediating anxiety-like behavior (5). Repeated activation of these pathways reinforces rewarding effects and contextual cue-drug memories that contribute to craving and relapse vulnerability (6).

Currently pharmacological approaches for smoking cessation include nicotine replacement therapy, varenicline, and bupropion (7). Although these treatments reduce withdrawal symptoms and craving by modulating dopaminergic signaling (8), their effectiveness remains limited due to side effects, low adherence, and high relapse rates. Hence the need for new pharmacological approaches for inhibiting the rewarding effects underlying nicotine use.

Aripiprazole is an antipsychotic drug acting as a partial agonist at the dopamine D_2_ receptor. This pharmacological profile allows it to act as a dopamine stabilizer during excessive peaks or downs, such as those induced by drugs of abuse(9). Importantly, recent evidence has pointed to a role for serotoninergic neurotransmission in nicotine addiction (10), therefore, aripiprazole serotoninergic mechanism of action as a 5-HT_1A_ agonist and 5-HT_2A_ antagonist may contribute to its therapeutic potential. Preclinical and clinical studies suggested that aripiprazole can reduce craving and drug-seeking behaviors associated with alcohol and cocaine although its efficacy remains controversial (11–15).

Animal models of anxiety, such as the elevated plus maze (EPM), and place conditioning paradigms, namely conditioned place preference (CPP) and conditioned place aversion (CPA), are widely used to investigate the effects of drugs of abuse. The EPM provides useful insights in the role of anxiolytic-like response in drug abuse, whereas CPP and CPA protocols mimic the contextual memory and cue-associated components of drug reward and aversion in experimental animals (16). Nicotine often produces inconsistent findings compared to other drugs of abuse in both the EPM (17) and place conditioning (18) protocols. The literature reports either the induction of anxiolytic-like effects and CPP or anxiogenic-like responses and CPA, reflecting its dose-dependent effects in humans. In addition, female subjects remain underrepresented in preclinical studies, despite evidence that sex differences influence nicotine sensitivity, reward processing, and relapse vulnerability (19,20).

Here, we combined behavioral assessments to investigate the aversive and rewarding properties of nicotine in male and female mice and tested the hypothesis that aripiprazole prevents nicotine-induced rewarding effects.

## Materials and Methods

### Animals

Male and female C57BL/6J wild-type mice (20–25 g) with 8-9 weeks of age were housed in groups of ten animals per box in a room under constant temperature (24 ±2 °C) and a 12 h light/dark cycle, with free access to water and food. The experiments were conducted during the light phase of the cycle. All procedures were approved by the Institutional Animal Care and Use Committee of the Federal University of Minas Gerais (CEUA/UFMG) under protocol number 256/2022, according to Brazilian and international guidelines for the use of laboratory animals.

### Drugs

Nicotine hydrogen tartrate salt (0.25,0.5 and 1 mg/kg; Sigma Laboratory) was diluted in saline (NaCl 0,9%) with pH normalized to physiological levels and administered via subcutaneous route. Aripiprazole (0.1, 1 and 10 mg/kg; Brystol-Myers Squibb / Otsuka Pharmaceuticals) was dissolved in a vehicle consisting of Cremophor/Ethanol/Saline (1:1:18) and administered via intraperitoneal route. Both drugs were administered at a volume of 10 ml/kg. The dose ranges were selected based on previous studies (21).

### Behavioural assessments

#### Locomotion in the open field

Locomotor activity was measured in an open field, consisting of a circular arena with 30 cm diameter and 50 cm height of opaque acrylic walls and a white floor (22). Animals received nicotine administration (0.25, 0.5 and 1 mg/kg, s.c.) and were immediately placed in the arena for a 20-min test session. The doses of nicotine were based on previous studies (21). Total distance travelled was recorded and analysed using AnyMaze software (Stoelting Co®). The apparatus was cleaned with 30% ethanol between animals.

#### Elevated plus maze (EPM)

The apparatus consisted of two opposite open arms (30 x 5) crossed perpendicularly by two arms of the same dimensions, enclosed by 40 cm walls height. The maze was elevated 50 cm above the floor, and all the experiments were carried out with lighting at 100 lux. Animals received nicotine administration (0.25,0.5 and 1 mg/kg, s.c.) and were immediately placed in the centre of the maze facing an open arm and freely explored the apparatus for 5 minutes. Anxiety-like behavior was assessed by measuring the percentage of time spent in the open arms and percentage of open arm entries (excluding time and entries in the central zone). Locomotor activity was evaluated by the absolute number of closed arm entries, which provides an index of general exploratory activity. All parameters were recorded and analysed using ANY-maze software (Stoelting Co®) (23).

#### Conditioned place preference (CPP)

As previously described (21)nicotine-induced CPP was assessed using a two-compartment acrylic apparatus (15×12×12 cm each) connected by a central corridor (9.5×5×12 cm) with removable doors. The two compartments differed in visual and tactile cues (distinct wall patterns and floor textures). On the first day (pre-test or trial 1; day 1), each animal was placed in the central compartment with free access to the entire box for 15 min to assess baseline preference. Animals spending more than 70% of the time in one of the compartments were excluded from the study. During the conditioning phases (days 2–7), animals received vehicle injections and were confined to one compartment for 30 min on days. On days, animals received nicotine (0.25, 0.5 and 1 mg/kg) immediately before being confined to the opposite compartment for 30 min. The doses of nicotine were based on previous studies (21). On day 8 (test day or trial 2), animals were again placed in the central compartment with free access to both compartments for 15 min, and time spent in each compartment was recorded. The apparatus was cleaned with 30% ethanol between animals.

To evaluate effects on the acquisition of nicotine-induced CPP, aripiprazole (0.1, 1 and 10 mg/kg) was administered 30 min before each nicotine injection. To test for its effects upon the expression of nicotine-induced CPP, aripiprazole (0.1, 1 and 10 mg/kg) was administered 30 min before exposure only on the test day.

The experiments were recorded with a video camera (Microsoft LifeCam®) and the time spent in each compartment was analysed using ANYmaze software (8ª version). CPP score (%) was calculated as: (time in drug-paired compartment − time in vehicle-paired compartment)×100/ (total time in both compartments). Data were graphically represented in two ways: (1) as the percentage mean of time spent in the drug-paired compartment during the pre-test (PT) and test (T) sessions, and (2) as individual test-day values to illustrate data distribution.

### Statistical analysis

The sample size appropriate for each experiment was estimated based on pilot studies. The animals were randomized for experimental treatments. The data were analysed for normality with Shapiro-Wilk test and the results using one or two-way analysis of variance (ANOVA), considering treatment (drug factor) and trial (pre-test vs. test in the CPP protocol) as experimental factors, followed by Tukey post hoc test. The significance level was considered at p < 0.05. The data are expressed as the mean and standard error of the mean (S.E.M.) superimposed to individual values. GraphPad Prism (9 version) was used to analyse the data and generate the graphs.

**Fig 1:**
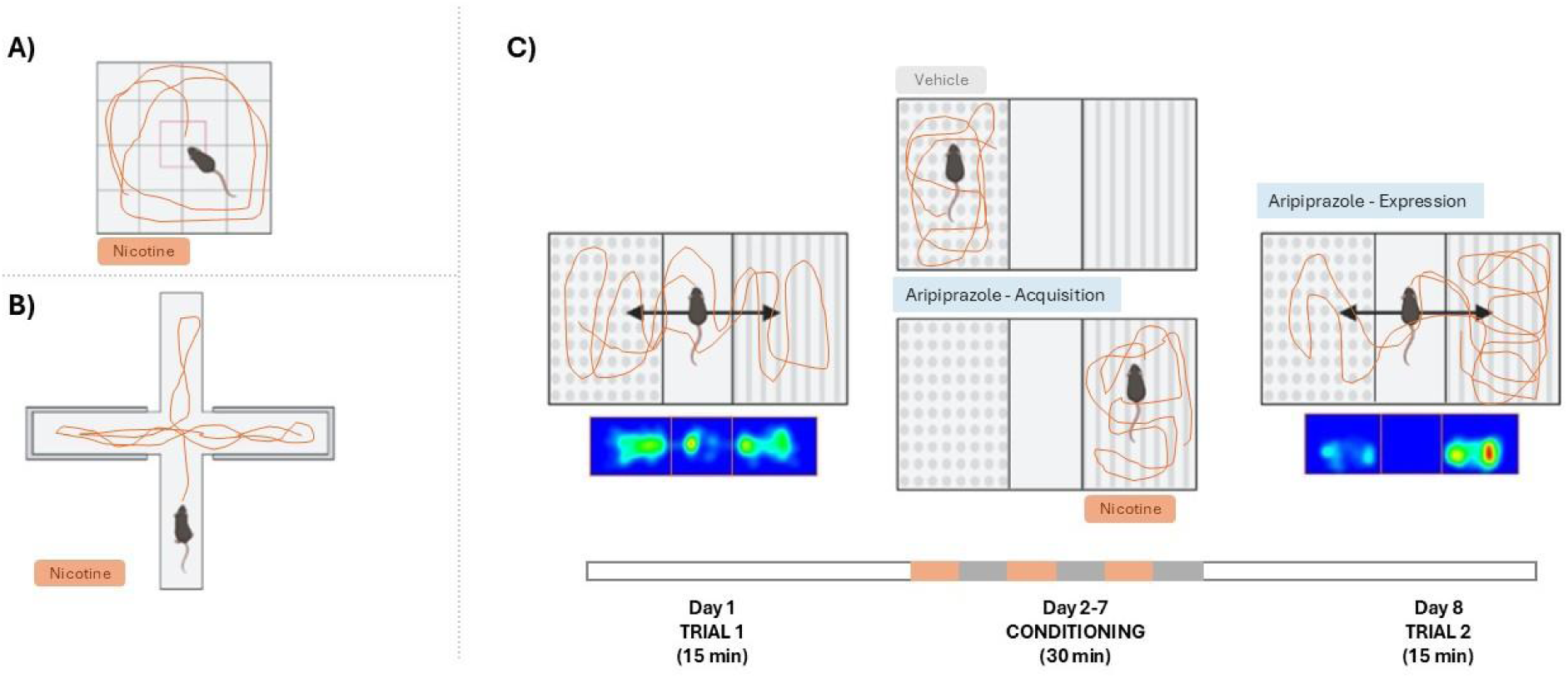
Experimental design. A) Open Field. B) Elevated Plus Maze. C) Conditioned Place Preference.

## Results

### Nicotine (1 mg/kg) reduced basal locomotion in males and females in the open field

To evaluate the effects of nicotine on basal locomotor activity, total locomotion was quantified in the open field. One-way ANOVA followed by Tukey’s post hoc test revealed a significant effect of treatment in both males [F(3,36) = 11.40, p < 0.0001; Fig.2A) and females [F(3,34) = 6.60, p = 0.0012; Fig.2B]. In both sexes, nicotine at the highest dose (1 mg/kg) reduced locomotion compared to the control group.

**Fig 2:**
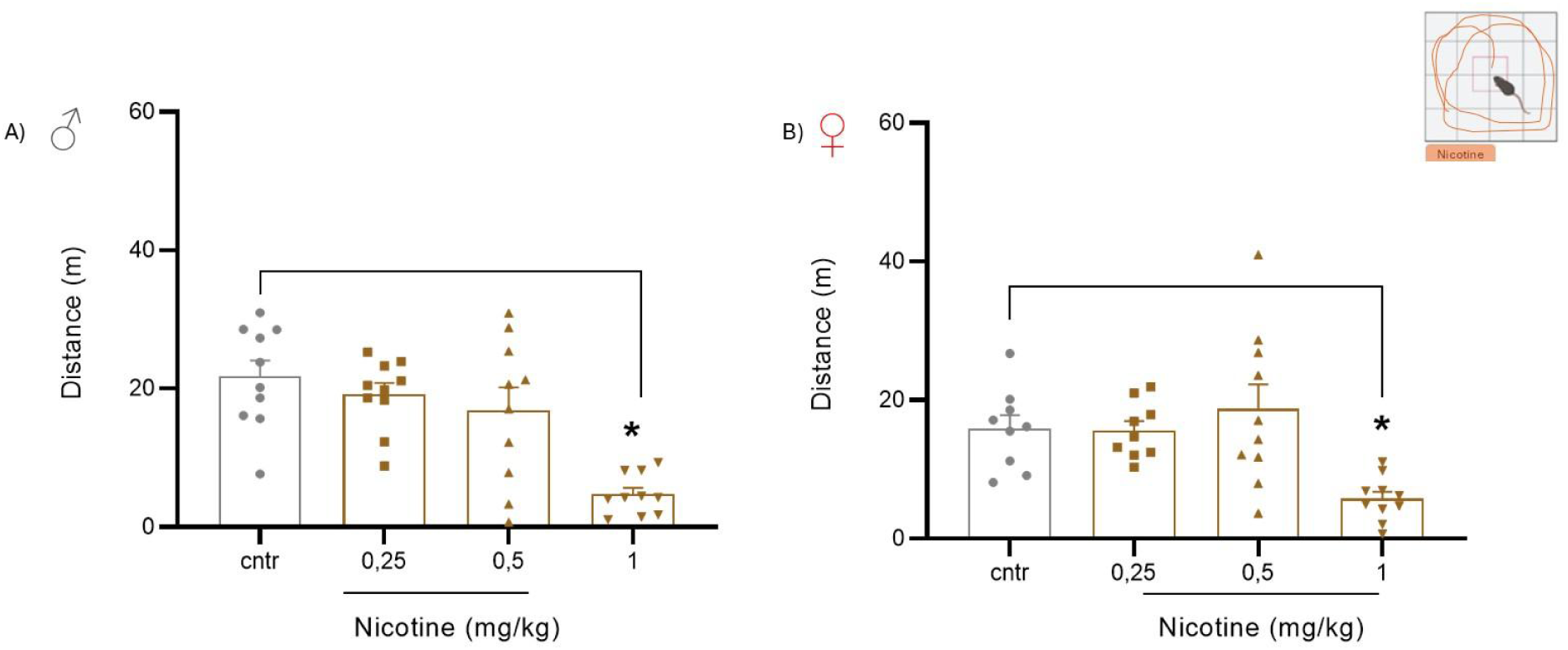
Effects of nicotine (0.25, 0.5 and 1 mg/kg) in the open field test (OFT) in A) males and B) females. Data are presented as mean ± SEM (n = 9–10 per group). One-way ANOVA followed by Tukey’s post hoc test revealed a significant reduction in locomotion at the highest dose (1 mg/kg) in both sexes (p < 0.05 vs. control).

### Nicotine (1 mg/kg) decreased open arm exploration in males in the EPM

In males, nicotine (1 mg/kg) decreased the percentage of time spent in the open arms in the elevated plus maze (EPM), as detected by one-way ANOVA [F(3,27) = 3.29, p = 0.0358; Fig.3A]. A similar effect was observed for the percentage of time spent in the enclosed arms [F(3,27) = 3.29, p = 0.0358; Fig.3B]. No significant differences were found for the percentage of open arm entries [F(3,27) = 1.89, p = 0.1551; Fig.3C] or absolute enclosed arm entries [F(3,27) = 1.7, p = 0.19; Fig.3D).

**Fig 3:**
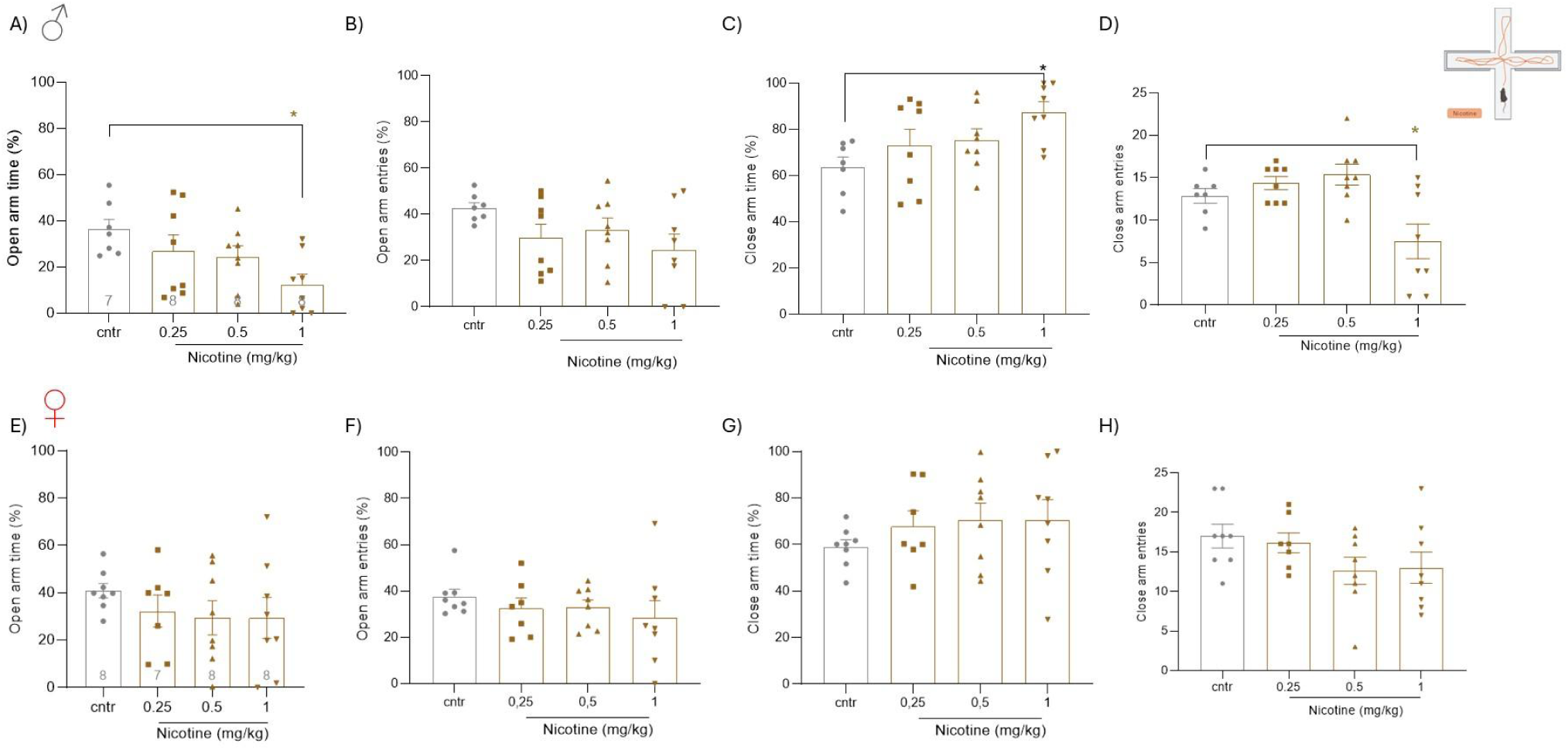
Effects of nicotine (0.25, 0.5 and 1 mg/kg) in the elevated plus maze (EPM) in A-D) males and E-H) females. Data are presented as mean ± SEM (n = 7–8 per group). One-way ANOVA revealed a significant effect of nicotine in males, with reduced percentage of time spent in the open and close arms and absolute close entries at the highest dose (1 mg/kg; p < 0.05, Tukey’s post hoc test), whereas no significant effects were observed in females.

In contrast, no significant effects of nicotine were observed in females in any EPM parameter, including percentage of open arm time [F(3,27) = 0.68, p = 0.5725; Fig.3E], percentage of open arm entries [F(3,27) = 0.61, p = 0.6118; Fig.3F], percentage of enclosed arm time [F(3,27) = 0.68, p = 0.5725; Fig.3G], or absolute enclosed arm entries [F(3,27) = 0.81, p = 0.75; Fig.3H).

### Nicotine (0.5 mg/kg) induced conditioned place preference (CPP) in males

In males, nicotine (0.5 mg/kg) increased preference for the drug-paired compartment (Fig.4A). One-way ANOVA showed significant drug effects on the test day, in which nicotine 0.5 mg/kg increased CPP score [F(3, 36) = 2.585, p = 0.06; Fig 4].

**Figure 4.**
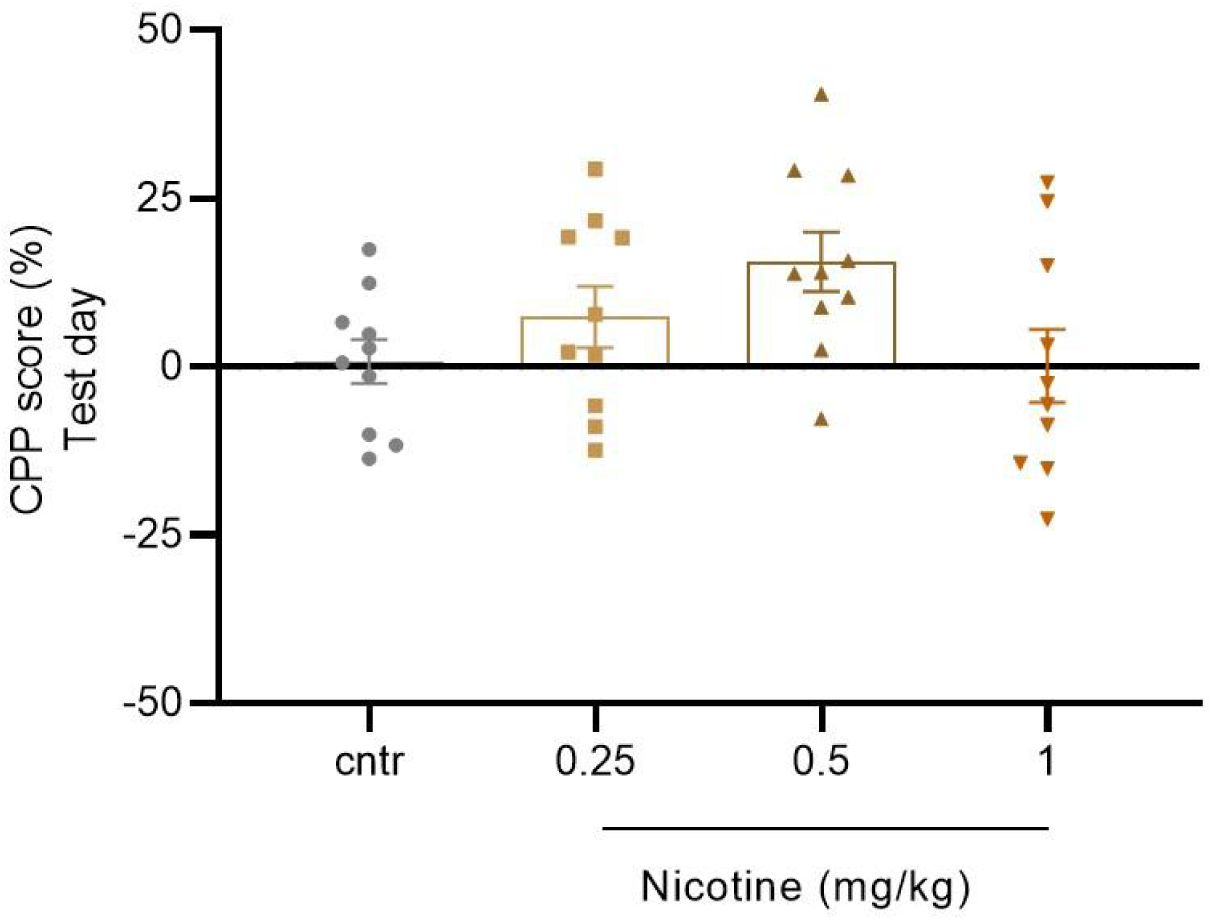
A) Nicotine induced CPP in male mice (0.5 mg/kg). Data are expressed as mean ± sem (left panel), as well as individual values (right panel), analysed by Two or One way ANOVA (n = 8-10).

### Aripiprazole prevented CPP acquisition in males

In accordance with our working hypothesis, aripiprazole prevented the acquisition of CPP when it was administered 30 min before each nicotine pairing in the CPP apparatus (Fig 5A). Two-way repeated measures ANOVA revealed no overall trial effect [F(1, 35) = 3.026. p = 0.0907] but a significant effect of treatment [F(3,35) = 8.48, p = 0.0002] and treatment × trial interaction [F(3, 35) = 3.768; p = 0.0192]. Consistently, analysis of the test session using one-way ANOVA confirmed a significant effect of treatment [F(3,35) = 18.24, p < 0.0001; Fig 5B]. Aripiprazole prevented CPP acquisition at all tested doses in males.

**Figure 5.**
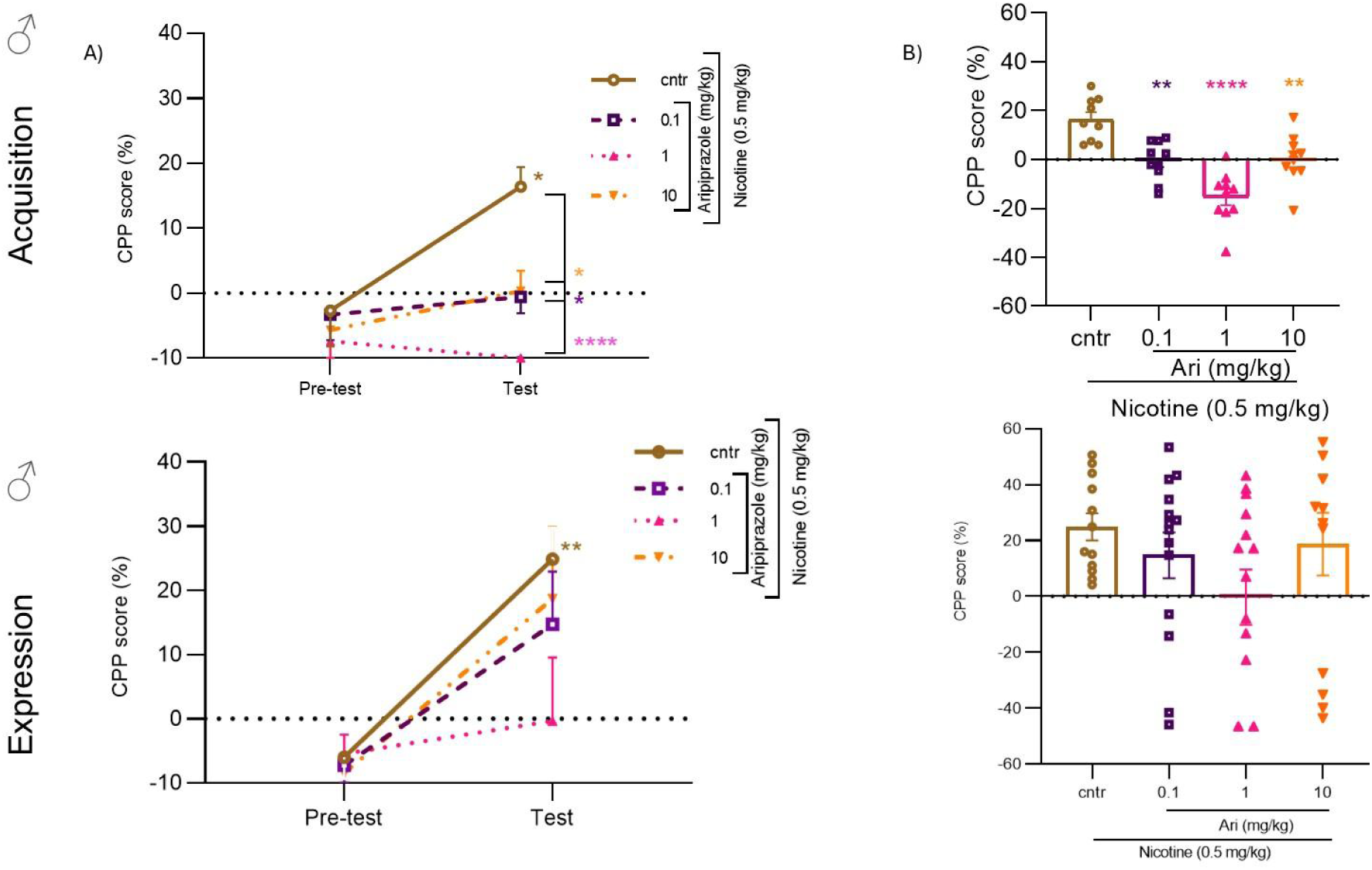
Aripiprazole inhibited nicotine-induced CPP in all doses during acquisition in males and on the test day with middle dose (1 mg/kg) A) Pre-test and post-test results. B) Test day with individual values. Data are expressed as mean ± sem (left panel), as well as individual values (right panel), analysed by One way ANOVA followed by Dunnet test (n = 14-16).

When aripiprazole was administered exclusively prior to the test session, it failed to interfere with CPP expression (Fig 5C). One-way repeated measures ANOVA showed no significant trial effect [F(1, 49) = 23.48; p < 0.0001], drug effect [F(3,49) = 0.93, p = 0.4340] or drug-trial Interaction [F(3, 49) = 1.699; p = 0.1796]. Moreover, one-way ANOVA of the test session showed no difference between groups [F(3, 49) = 1.424; p = 0.2470; Fig. 5D].

## Discussion

In the present study we investigated the behavioral effects of nicotine using a range of doses in male and female mice. At the highest dose (1 mg/kg), nicotine induced hypolocomotion in both sexes and reduced exploration in the EPM in males. The intermediate dose (0.5 mg/kg) failed to interfere with anxiety-like behavior but induced CPP in males. Finally, in line with our working hypothesis, aripiprazole prevented nicotine-induced CPP in male mice.

At the highest dose, nicotine reduced open arm exploration in the EPM in males. This response could be interpreted as an anxiogenic-like effect. However, this dose also reduced basal locomotion in the open field and the number of open arms entries the EPM. Thus any conclusion on anxiety-like behavior at this dose should be viewed with caution due to locomotor-impairing effect (24,25). Importantly, our results aligned with previous literature showing either aversive or motor-impairing effects (24,26).

Nicotine-induced CPP has been more consistently demonstrated in male rodents, and our findings are in line with previous studies using similar dosing and protocols (21). In contrast, nicotine CPP in females, and sex differences in sensitivity to nicotine reward and aversion remain poorly understood. Emerging evidence suggests that females may exhibit distinct neurobiological and behavioral responses to nicotine (27), highlighting the importance of including both sexes in experimental designs (28).

Sex differences in nicotine studies were already reported (29). It was previously observed that nicotine induced-CPP in males but not in females (27,30), similar findings were also described in humans (31). Intriguingly, different phases of the estrus cycle don’t seem to affect female performance, although ovariectomised female rats did not show CPP suggesting sex differences for nicotine response (32). In females, nicotine effects seem to be age-dependent, with adolescent female mice being more sensitive to the rewarding effects of nicotine and adults to the aversive ones (30). Nevertheless, previous studies showed that higher doses of nicotine are able to induce significant CPP in females, especially in rats, with 0.4 mg/kg (30) and 0.6 mg/kg (27), but also in mice 0.4 mg/kg (33).

In addition to characterizing the effects of nicotine on place conditioning, our study shows for the first time that aripiprazole, an antipsychotic drug, is effective in preventing the acquisition of CPP to nicotine in male mice. This effect occurred at all doses tested, suggesting a robust effect of this atypical antipsychotic. As for CPP expression, the intermediate dose (1 mg/kg) seems to reduce nicotine effect, although statistical analyses failed to detect an effect, possibly due to the remarkable data variability. These findings support the hypothesis that aripiprazole modulates nicotine reward probably by stabilizing dopaminergic signaling and interfering with associative learning processes related to drug cues.

Aripiprazole has been developed as a partial agonist at the dopamine D_2_ receptor, primarily as an antipsychotic drug, primarily for the treatment of psychotic disorders, such as schizophrenia (34). The underlying rationale was to develop an alternative to conventional antipsychotics and therefore to avoid the motor side effects (parkinsonism) resulting from dopamine antagonism (34). Considering the well established role of aripiprazole on regulating dopaminergic rewarding pathways, it seems reasonable to consider repurposing this drug and other dopaminergic modulators as pharmacological approaches to substance use disorders (35) preclinical models for studying psychostimulants, rewarding and reinforcing responses to various drugs of abuse. Aripiprazole prevented the effects of psychostimulant drugs, such as cocaine and amphetamine (36–38). Moreover, it inhibited CPP induced by morphine without interfering with its analgesic effects (39,40). Aripiprazole was also effective in preventing various abuse-related behavioural effects induced by ethanol administration (41,42). The present studies advances the literature by showing that aripiprazole also prevents the rewarding effects of nicotine.

In conclusion, the present finding demonstrates that dose and sex as important variables to be considered when investigating the anxiolytic and rewarding effects of nicotine in experimental animals. Nicotine effects on anxiety-like behavior were confounded by reduction in basal exploratory activity, whereas rewarding effects in the CPP were consistent only in male mice. Finally, our studies show that aripiprazole remarkably prevents nicotine acquisition of CPP in male mice. This antipsychotic drug may have therapeutic potential for smoking cessation, particularly during stages in which conditioned responses contribute to the maintenance of smoking behavior.

## Authors contributions

All authors contributed to the study conception and design. Material preparation, data collection and analysis were performed by Rayssa Brianis, Lucas Bedeschi. Lia Iglesias, Samia Joca, Daniele Aguiar, Fabrício Moreira provided mentoring during the whole project. The first draft of the manuscript was written by Rayssa Brianis. All authors commented on previous versions of the manuscript. All authors read and approved the final manuscript.

## Acknowledgements and funding

CNPq for their Research Productivity Fellowships (protocol 305181/2023-8 for FAR). The authors acknowledge CNPq for research grants (protocols: 444403/2024-8, 405689/2025-0 and 446844/2025-0). Support was also provided by FAPESP (protocol: 2023/16182-2) and INCT Substâncias Psicoativas and INCT Saúde Mental Digital.

